# Weaker neuroligin 2 – neurexin 1β interaction tethers membranes and signal synaptogenesis through clustering

**DOI:** 10.1101/2024.10.16.618688

**Authors:** Robbie Boyd, Khuloud Jaqaman, Weiwei Wang

## Abstract

Single-pass transmembrane proteins neuroligin (NL) and neurexin (NRX) constitute a pair of synaptic adhesion molecules (SAMs) that are essential for the formation of functional synapses. Binding affinities vary by ∼ 1000 folds between arrays of NL and NRX subtypes, which contribute to chemical and spatial specificities. Current structures are obtained with truncated extracellular domains of NL and NRX and are limited to the higher-affinity NL1/4-NRX complexes. How NL-NRX interaction leads to functional synapses remains unknown. Here we report structures of full-length NL2 alone, and in complex with NRX1β in several conformations, which has the lowest affinity among major NL-NRX subtypes. We show how conformational flexibilities may help in adapting local membrane geometry, and reveal mechanisms underlying variations in NL-NRX affinities modulation. We further show that, despite lower affinity, NL2-NRX1β interaction alone is capable of tethering different lipid membranes in total reconstitution, and that NL2 and NRX1β cluster at inter-cellular junctions without the need of other synaptic components. In addition, NL2 combines with the master post-synaptic scaffolding protein gephyrin and clusters neurotransmitter receptors at cellular membrane. These findings suggest dual roles of NL2 - NRX1β interaction – both as mechanical tether, and as signaling receptors, to ensure correct spatial and chemical coordination between two cells to generate function synapses.

## Introduction

Neuronal cells transmit and process information through synapses, where the plasma membranes of the pre- and the post-synaptic neurons are tethered by synaptic adhesion molecules (SAMs)(*1, 2*). Post-synaptic protein neuroligin (NL) and pre-synaptic protein neurexin (NRX) constitute a pair of SAMs that play indispensable roles in synapse function(*1, 3, 4*).

Knockout of NL or NRX results in severe defects in synaptic transmission, with NL deletion being fatal for postnatal mice(*5–8*). Genetic mutations in humans have been associated with neuropsychiatric disorders including autism, intellectual disabilities, and epilepsy(*9–14*).

Although not seeming to change the number of synapse-like structures in some tissues, genetic ablation of NL and NRX cripples synapse function throughout the nervous system (*2, 15–17*).

Multiple NL and NRX subtypes are found in mammalian nervous systems. NL has four major subtypes, NL1-4, that differ in tissue distribution and binding affinities with NRX. NL1 and NL2 preferentially localize in excitatory and inhibitory synapses, respectively, while NL3 and NL4 show less/unclear selectivity(*11, 18–21*). NL1 and NL4 bind NRX with nanomolar to sub-micromolar affinities, while NL2 and NL3 bind weaker in the micromolar range(*22, 23*). NRX family contains major three genes, NRX1-3. Each has a long “α” form containing 6 extracellular “laminin, neurexin and sex hormone – binding globulin–like” (LNS) domains, and a short “β” form containing only the last LNS domain that binds with NL. Combined with splice variants, more than a thousand subtypes are possible for NRX that differ in affinities for NL and other ligands. The wide range of NL-NRX interaction affinities has been implicated in chemical specificity of synapses(*8, 24–26*).

A few fundamental questions remain unanswered regarding how NL – NRX interaction works at synapses. NL and NRX are both single-pass transmembrane proteins. Structural information to date is limited to truncated (or possibly degraded) NL and NRX containing only the extracellular acetylcholinesterase (AChE) domain for NL, and the LNS domain for NRX(*10, 23, 27–33*), limiting understanding of the full-length conformation. In addition, despite higher affinity complex structures (NL1-NRX and NL4-NRX) having been resolved almost 20 years ago(*27, 28*), lower affinity complexes (NL2/3-NRX) have not been reported. This has obscured the mechanism underlying lower affinities and put into question whether low affinity allow stable complex and tether of cellular membranes.

To address the above questions, we resolved structures of full-length NL2 alone, and in complex with full-length NRX1β. These structures reflected flexible orientations between extracellular domains (ECDs) and transmembrane domains (TMs) for both NL and NRX, as well as within dimeric NL2 ECD. The mechanism underlying low NL2-NRX1β differ from that previously proposed(*29*). We further show that NL2 and NRX1β are clustered at inter-cellular junctions in non-neuronal cells, and that their interaction alone is capable of tethering two lipid membranes, suggesting a signaling role leading to the formation of synapse. Co-clustering of NL2, gephyrin and glycine receptor at the cellular membrane suggests roles in recruiting neurotransmitter receptors to the synapse, and in signaling pre-synaptic cell through clustering of NRX. A mechanism through which weaker NL2-NRX1β interaction both tethers membranes and signals the recruitment of other synaptic components through clustering, has been proposed.

## Results

### Flexible linker between ECD and TM

In full-length neuroligins, the Ser- and Thr-rich linker (∼ 60 aa) between the ECD (∼550 aa) and the transmembrane helix (∼40 aa, TM) is named the stalk domain since it contains glycosylation sites and may provide some structural rigidity and spacing between the ECD and TM(*23, 29, 31, 34*) (Fig. 1A). The intracellular domain (ICD, ∼130aa) that interacts with multiple synaptic proteins is believed to be unstructured. Reported structures of neuroligins are determined using truncated proteins containing only the extracellular domain (ECD)(*10, 27–33*). To examine whether the ECD has defined/preferred orientations with respect to the stalk/TM/ICD(*23*), we analyzed full-length NL2 using single particle cryo-EM.

**Figure 1.**
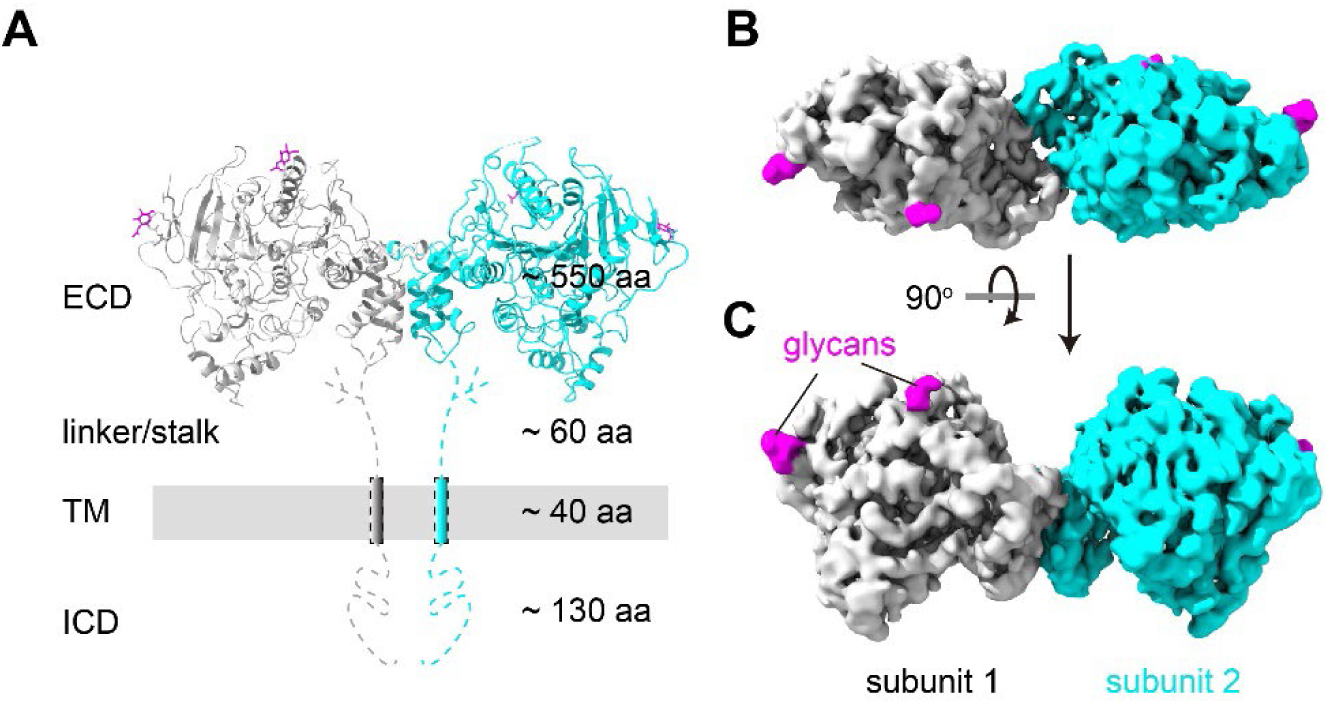
Structure of full-length NL2 contained only the extracellular domain. (**A**) Schematic of full-length NL2. Grey bar denotes post-synaptic membrane. Cryo-EM density map of NL2 viewed (**B**) down extracellular side and (**C**) parallel to membrane. Two subunits are colored in grey and cyan, respectively. Glycans are colored in purple.

EM densities for the stalk, TM (detergent micelle) and intracellular domains are mostly absent for full-length NL2 (Fig. S1A, B, E and Fig. S2 A, B). Only the ECD was well resolved, at an overall resolution of 3.3 Å (Fig. 1B and Fig. S2 C-F), allowing for unambiguous model building (Fig. 1A, see methods for details). Two AChE domains of NL2 form a dimer, like that observed in ECD-only structures(*10, 11, 27, 28, 32, 33, 35*). Apparently, the stalk domain provides very limited rigidity between ECD and TM.

### Full- and sub-stoichiometric complexes of NL2-NRX1 with structural flexibility

Among neuroligin subtypes NL1-NL4, NL2 has the lowest apparent affinity with NRX1β(*22, 23*). Complex structures of NL1-NRX1β (*K_d_* ∼ 10 nm) and NL4-NRX1β (*K_d_* ∼ 100 nm) ECDs have been resolved nearly 2 decades ago(*23, 27, 28*). NL3 binds NRX1β with ∼ 1 µM *K_d_* and NL2 binds even weaker at ∼ 10 µM *K_d_*(*22, 23*). So far, no NL2/NL3-NRX complex structure has been reported. Whether or not the weaker NL2/3-NRX1 interactions enable stable complex has not been established.

Mixing of purified full-length NL2 and NRX1β led to co-eluting peak in size exclusion chromatography (Fig. S1B-D). Like NL2 alone, single-particle cryo-EM analysis resolved only the ECDs of NRX1β and NL2 (Fig. S2G-M), suggesting non-rigid connection between ECDs and TMs for both proteins (Fig. 2A). Three conformations of full stoichiometric (2:1 NRX1β:NL2 dimer) complex, differing in relative orientations between two subunits of NL2 dimer, were resolved at 3.2 Å overall resolution (Fig. 2B, C, Fig. S2K-M Conf. 1-3). One sub-stoichiometric complex (1:1 NRX1β:NL2 dimer) was also resolved at 3.9 Å overall resolution (Fig. 2D, E, Fig. S2K-M Conf. 0).

**Figure 2.**
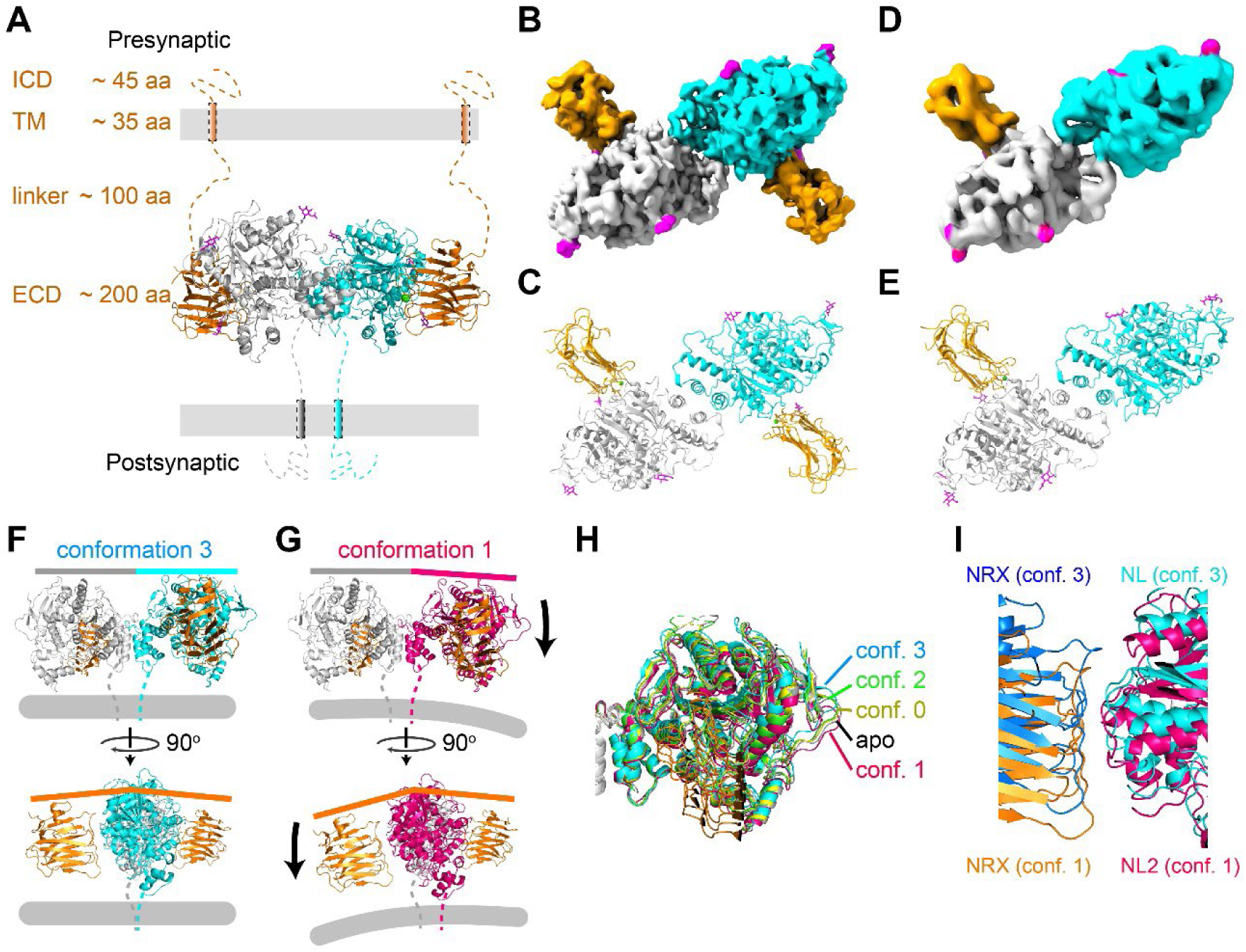
Overall conformations of full- and sub-stoichiometric NL2-NRX1β complexes. (**A**) Illustration of NL2-NRX1β complex at synapse. Two NL2 subunits are colored in grey and cyan, respectively. NRX1β and glycans are colored in orange and purple, respectively. (**B**) Density map and (**C**) atomic model of a 2:1 NRX1β:NL2 dimer full stoichiometric complex. (**D**) Density map and (**E**) atomic model of the 1:1 NRX1β:NL2 dimer sub stoichiometric complex. Different orientations between two NL2 subunits in (**F**) conformation 3 and in (**G**) conformation 1. Grey bars indicate direction of post-synaptic membrane. (**H**) Comparison of one NL2 subunit orientations in all structures, with the other subunit aligned (grey: aligned subunit). Color codes shown. (**I**) NRX1β and NL2 moves together and preserve interaction interface.

NL2 dimer showed structural flexibility between its two monomers. Comparing the three structures of full stoichiometric complexes (Fig. S2M, conf. 1-3), a pivoting motion between two monomers becomes evident. This motion led to change in orientation between two monomers when viewed in parallel to the membrane (Fig. 2F, G). The sub stoichiometric complex (conf. 0) had very similar orientation as apo NL2, both sitting roughly in the center of the range observed in full stoichiometric complexes (Fig. 2H). NL2 and NRX1β move in coordination in all complex structures and maintain an unchanged interaction interface (Fig. 2I). Since the binding of NRX1β does not result in unidirectional change in NL2 monomer orientations, the pivoting motion is likely arising from intrinsic NL2 flexibility.

### Mechanisms underlying weak NL2-NRX1β interaction

The NL2-NRX1β interaction interface resembles that of NL1-NRX1β, composed of mostly polar interactions with a small buried-area of ∼ 1,000 Å^2^ (1,160 Å^2^ for NL1-NRX1β) with a Ca^2+^ mediating multiple interactions (Fig. 3A, B). NRX1β uses an essentially identical binding site to form NL1- and NL2-complexes. Ca^2+^ mediated interactions are also strictly conserved. However, two unique features of NL2 likely contribute to its much lower affinity.

**Figure 3.**
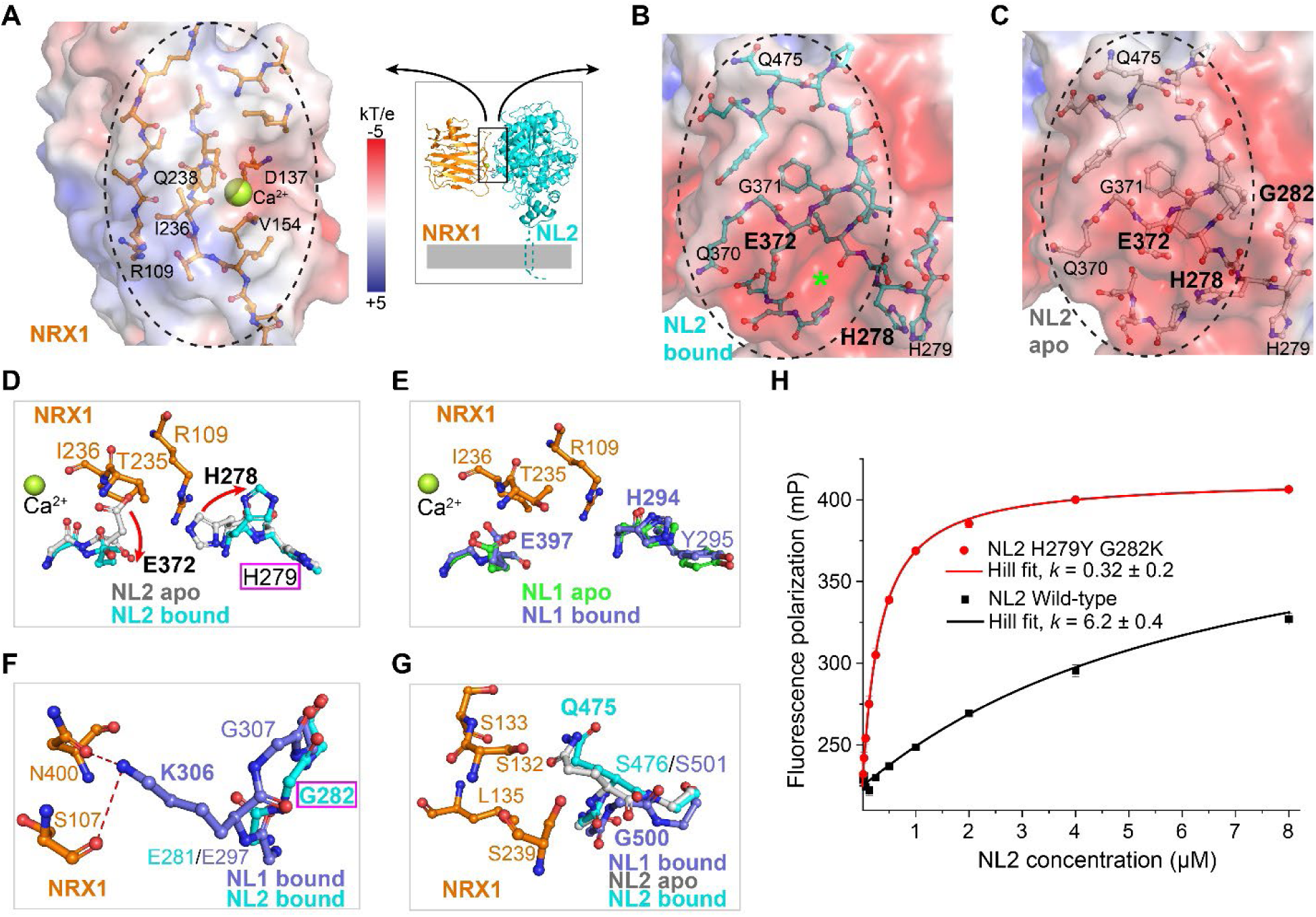
Structural features contributing to lower NL2-NRX1β affinity. Binding interfaces of (**A**) NRX1β, (**B**) NL2 bound and (**C**) NL2 apo. Molecular surfaces are colored according to electrostatics calculated using APBS. Amino-acid residues at the interfaces are shown as sticks. Green star indicates cavity in bound NL2 that interact with NRX1β:R109, which is buried in apo NL2. (**D**) Conformational changes of E372 and H278 between bound and apo NL2 open or occlude NRX1β:R109 binding cavity. (**E**) in NL1, the NRX1β:R109 binding cavity is present in both apo- and bound states. (**F**) NL1:K306 makes polar contacts with NRX1, but is mutated to G282 in NL2. (**G**) NL1:G500 mutation to NL2:Q475 does obstruct NRX1 binding or change conformation from apo to bound states. (**H**) Fluorescence polarization of NRX1 upon binding with wild-type NL2 (black) or H279Y G282K mutant (red), fitted with Hill equation with *k* values indicated and shared slope *n* = 1.05 ± 0.05.

First, interaction with NRX1β requires binding site rearrangements in NL2, but not NL1 (Fig. 2B-D). R109 of NRX1β binds to a negatively charged cavity on NL2 between E372 and H278 (Fig. 3B, green star). However, in apo state NL2, E372 and H278 sidechains occupy this cavity, preventing NRX1β:R109 from binding (Fig. 3D). In contrast, this cavity is available and does not change between the apo and bound states of NL1(*27, 29*) (Fig. 3E). Pre-positioned binding site for NRX1β most likely contributes to the higher affinity of NL1 than NL2.

Second, a non-conserved residue, NL2:G282 v.s. NL1:K306, provides less polar interactions between NRX1β and NL2 (Fig. 3F), thereby contributing to lower NL2 affinity. We note that the non-conserved residue NL2:Q475 (G500 in NL1) does not change conformation from apo to bound states (Fig. 5G). Therefore, the previously proposed mechanism of lower NL2 affinity due to Q475 steric hindrance is unlikely(*29*).

Non-cooperative binding may be an additional contributor. In the sub-stoichiometric structure (Fig. 2D, E, Fig. S2M, conf. 0), the bound and the unbound NL2 subunits exhibited different conformations, respectively corresponding to the bound and apo states. This suggests a lack of cooperativity between the NRX1β binding sites of the two NL2 monomers, consistent with reported binding isotherms following single-site binding models(*27, 29*). Being non-cooperative prevents increased affinity through multiple binding events on the same NL2 dimer.

Mutations at the above two interaction sites of NL2 dramatically improved affinity for NRX1β, mimicking NL1. In analysis of why NL2:H278 adopted a different configuration than the corresponding NL1:H294 (Fig. 3D, E), we found that the adjacent residue is not conserved, namely H279 in NL2 and Y295 in NL1. We reason that in NL2, H278 and H279 will carry the same charge and cause repulsion, while in NL1 H294 and Y295 does not repel each other, leading to the difference in NL2:H278 / NL1:H294 conformations in the apo state. A H279Y mutation in NL2, combined with G282K mutation to mimic NL1 (Fig. 3D, F purple boxes), dramatically increased apparent NRX1β affinity from ∼ 6 µM to ∼ 0.3 µM (Fig. 3H). Cleary these sites play major roles in the modulation of NL-NRX affinities.

### NRX1β and NL2 interaction is sufficient for tethering lipid membranes

In each synapse, there are multiple adhesion proteins that contribute to tethering of the two cellular membranes. Since NL2 - NRX1β interaction is the weakest among all NL-NRX pairs, it is unclear whether the NL2 - NRX1β interaction alone is sufficient for tethering lipid membranes. To address this without complications from undefined cellular environment, we employed a total reconstitution system (Fig. 4).

**Figure 4.**
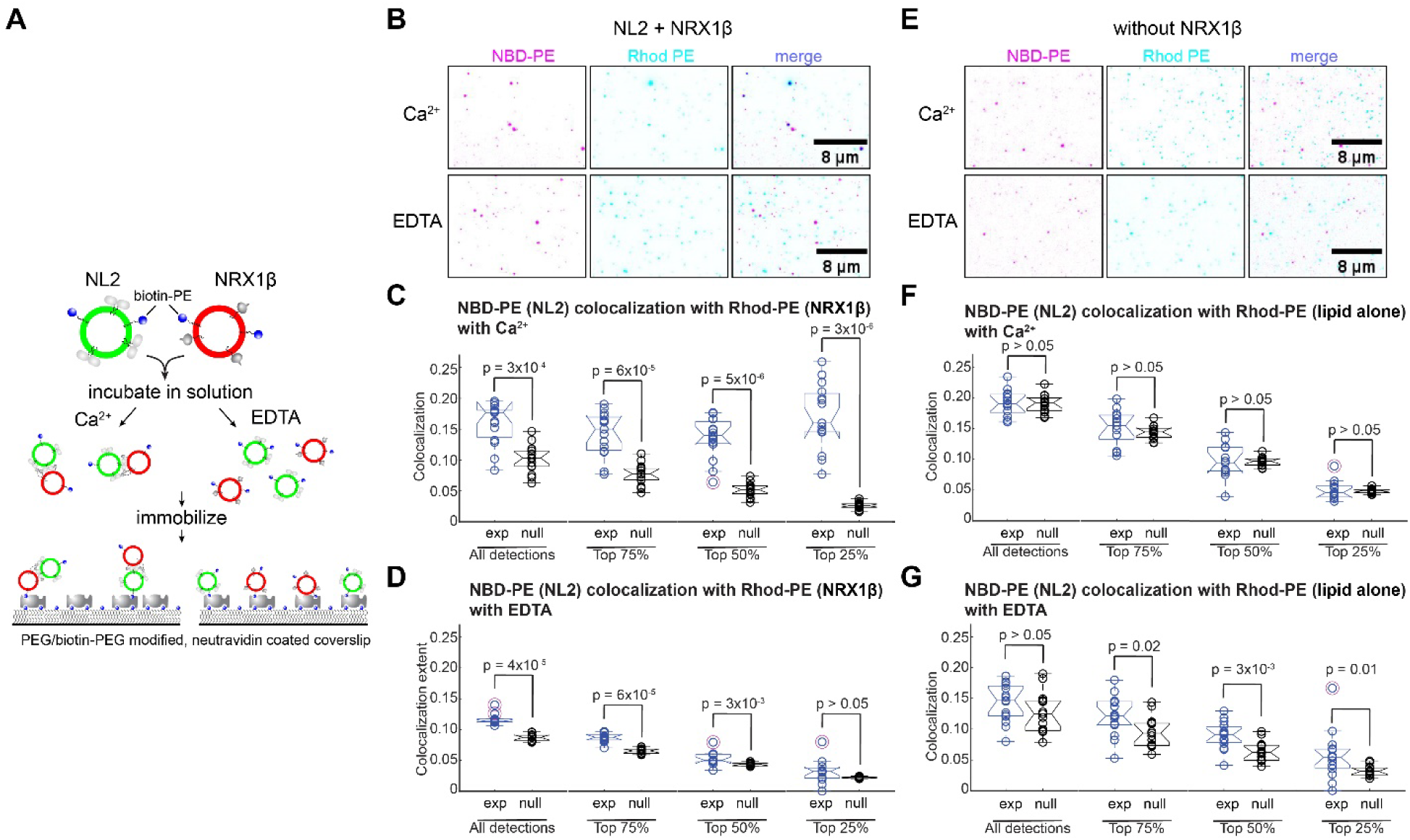
NRX1β and NL2 interaction tethers lipid membranes in reconstituted proteo-liposomes. (**A**) Schematic of the vesicle colocalization assay. Non-labeled wild type NL2 and NRX1β were used. Fluorescence originates from NBD-PE (green) and Rhodamine-PE (Red) in lipid vesicles. (**B**) Representative micrographs with inverted contrast of NL2 (NBD-PE) and NRX1β (Rhod-PE) vesicles in the presence (top panels) and absence (bottom panels) of Ca^2+^. Object-based co-localization analysis of NL2 (NBD-PE) and NRX1β (Rhod-PE) vesicles in the presence of (**C**) Ca^2+^ or (**D**) EDTA. (**E**) Representative micrographs of NL2 (NBD-PE) and lipid alone (Rhod-PE) vesicles in the presence (top panels) and absence (bottom panels) of Ca^2+^. Co-localization analysis of in the presence of (**F**) Ca^2+^ or (**G**) EDTA. In C, D, F, G, each circle represents one field of view, collectively summarized in a boxplot. For each box, the central mark is the median, the edges are the 25th and 75th percentiles, and the dashed whiskers extend to the most extreme inlier data points. Circles surrounded by an additional red circle indicate outlier datapoints (as deemed by the Matlab “boxplot” function). Notch around median indicates the 95% confidence interval of the median. Number of fields of view used in analysis = 15 (C), 12 (D), 13 (F) and 15 (G).

Full-length NL2 and NRX1β wild-type were reconstituted separately into proteoliposomes doped with biotinylated lipids, as well as green (NBD-PE, NL2) and red (Rhod-PE, NRX1β) fluorescent lipids (see methods for details). Reconstituted proteoliposomes were incubated in solution to allow for unobstructed interaction, and subsequently immobilization through biotin-neutravidin interaction on PEG-passivated glass coverslips (Fig. 4A). Total internal reflection microscopy (TIRF) was used to image immobilized vesicles, followed by distance-based colocalization analysis of detected vesicles(*36, 37*).

Ca^2+^ promotes colocalization of NL2 with NRX1β proteoliposomes in a manner dependent on vesicles size (largely reflected as detected object intensity, given the diffraction-limited nature of most of the vesicles) (Fig. 5B-D, Fig. S4A-C). Visually, more colocalization is apparent in the presence of Ca^2+^, compared with no Ca^2+^ (EDTA as chelator) (Fig. 4B). Interestingly, when analyzing all detected vesicles, the difference between with and without Ca^2+^ was unclear (Fig. 4C, D, “All detections”). However, when focusing on brighter and brighter detections (Fig. S4A), Ca^2+^-dependent colocalization becomes very significant while colocalization in its absence diminishes (Fig. 4C, D). Vesicles with the top 25% intensities (i.e. intensities ≥ 75^th^ percentile; Fig. S4A) showed clear, significant colocalization in the presence of Ca^2+^ with a *p* value ∼ 3 x 10^-6^ against the *computational null control* (see methods for details)(*36*), compared with *p* value > 0.05 without Ca^2+^ (Fig. 4C, D, “Top 25%”). Analysis of the converse colocalization of NRX1β with NL2 vesicles yielded similar outcomes (Fig. S4B, C). Since fluorescence is arising from fluorescent lipids at a given mole fraction, brighter detections represent larger vesicles within diffraction limit, which in turn contain more NL2 and NRX1β proteins on average for tethering membranes. Since Ca^2+^ binds at the interaction interface between NL2 and NRX1β (Fig. 3A) and increases binding affinity(*22, 27*), Ca^2+^ dependence further suggests a NL2-NRX1β interaction-dependent tethering.

**Figure 5.**
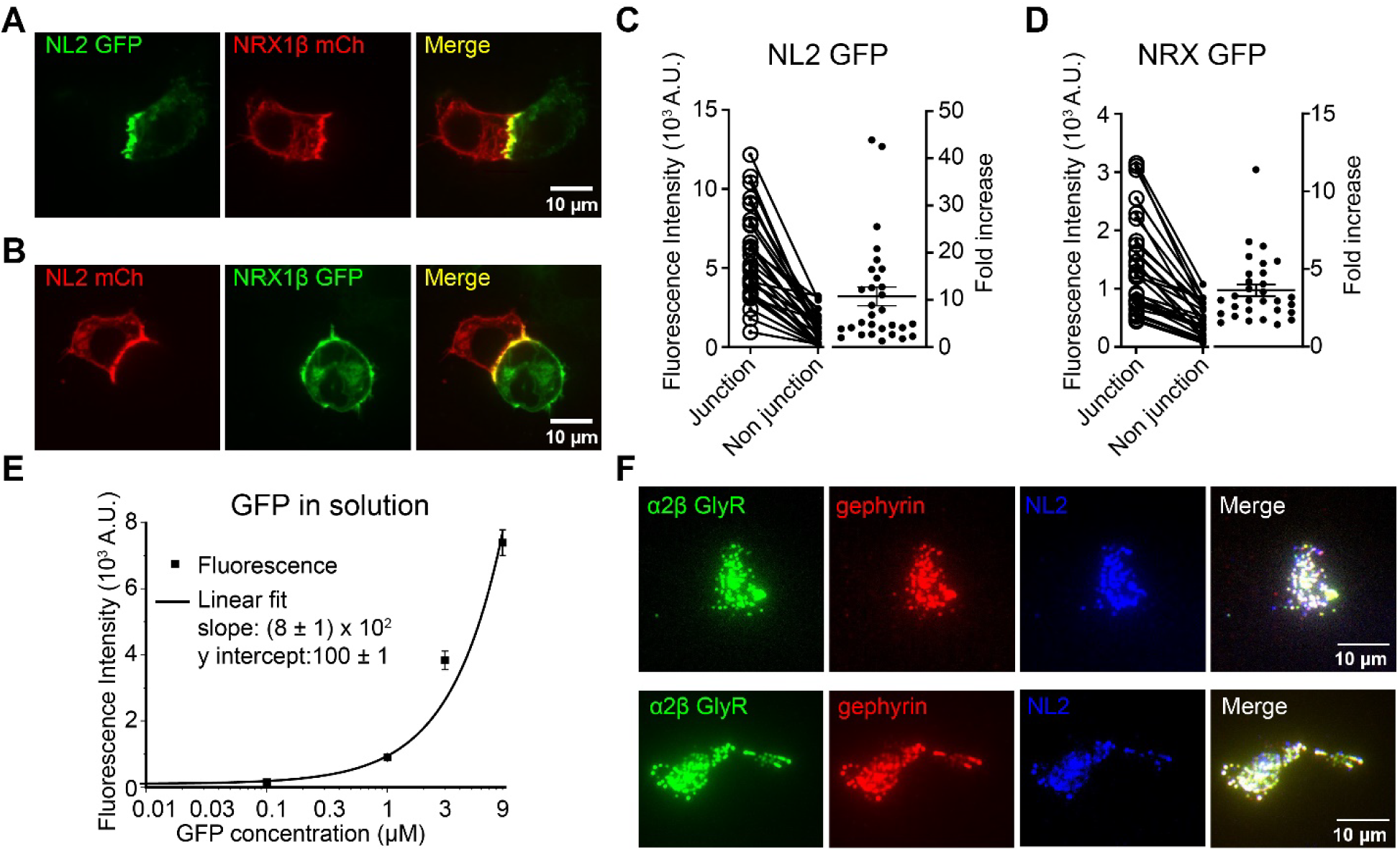
Quantification of NRX1β and NL2 at inter-cellular junctions. Representative confocal micrographs of (**A**) NL2-GFP and NRX1β-mCherry, (**B**) NL2-mCherry and NRX1β-GFP expressed in separated HEK293 cells and cultured together afterwards. Quantification of (**C**) NL2-GFP, (**D**) NRX1β-GFP fluorescence (n = 30 cells) at and outside of intercellular junctions. (**E**) Calibration of fluorescence intensity against GFP in solution. (**F**) Representative TIRF micrographs of unroofed cells expressing α2β GlyR (green), gephyrin (red) and NL2 (blue), with merged image shown.

Colocalization of vesicles requires the presence of both NL2 with NRX1β proteins. To evaluate protein-independent vesicle-vesicle interaction, we tested colocalization between NL2 proteoliposomes and lipid-alone vesicles that have the same lipid composition as NRX1β proteoliposomes, but without NRX1β (Fig. 4E-G). No colocalization was obvious with or without Ca^2+^ (Fig. 4E). Quantitative analysis shows no significant colocalization (when tested against *null* controls), with *p* values mostly greater than 0.01, across all vesicle sizes (Fig. 4F, G and Fig. S4D, E). Clearly, a NL2-NRX1β interaction-dependent membrane tethering was observed in reconstituted lipid vesicles.

### NRX1β and NL2 cluster at inter-cellular junctions

In neurons, NRXs and NLs are found clustered at opposing pre- and post-synaptic membranes(*8, 11*). It is unclear whether local concentrating of NRXs and NLs requires other components in the synapse, including other SAMs and intracellular scaffolding proteins and receptors. To test this, we expressed eGFP- and mCherry-fused NL2 and NRX1β in separate HEK293 cells and cultured them together. Apparently, NL2 and NRX1β are much more concentrated at inter-cellular contact sites (junctions) than other regions of the plasma-membrane, regardless of the fused fluorescent proteins (Fig. 5A, B). Since HEK293 cells are incapable of forming synaptic specializations, concentration of NL2 and NRX1β at cellular interfaces does not require the formation of synapse.

We quantified both NL2 and NRX1β in their eGFP-fused form (Fig. 5C, D) to avoid variables arising from differences in fluorescent protein fusions. Confocal slices were taken in the direction (largely) perpendicular to the cellular junctions. The fluorescence intensities increased an average of 10 ± 2 folds at junction for NL2 (Fig. 5C), and 3.6 ± 0.4 folds for NRX1β (Fig. 5D). To obtain estimates of NL2 and NRX1β concentrations, we calibrated the fluorescence intensities against eGFP concentrations in solution (Fig. 5E). The confocal volume was estimated based on the numerical aperture and wavelength (see methods for details), which should provide sufficient accuracy at the higher concentrations (> 100 nM) that we used(*38*).

The number of NL2 dimers per each µm^2^ of junction membrane is largely in the range of 500 to 2000, while that outside of junction is generally below 250. NRX1β is at lower concentrations, with ∼ 400 / µm^2^ at junctions and ∼ 100 outside.

Clustering of NL2 and NRX1β at inter-cellular junctions suggests a likely mechanism for bridging two membranes despite low affinities - multiple NL2 and NRX1β interactions provide sufficient avidity. In addition, since the intracellular portion of NL2 and NRX1β are known to bind with synaptic scaffolding proteins and organizers in the post- and pre-synaptic specializations, respectively(*8, 11, 26*), clustering of NL2 and NRX1β likely serves as a local signal to coordinate synaptogenesis between two neurons through recruiting correspondent synaptic components.

### NL2 co-clusters with gephyrin at the membrane and recruits α2β GlyR

NL2 is known to bind with the master scaffolding protein, gephyrin, which in turn binds with neurotransmitter receptors such as glycine and GABA receptors, as well as cytoskeleton and organizes functional inhibitory synapses(*11, 39, 40*). To evaluate whether NL2 clustering indeed signal the generation of functional synaptic specializations, we co-expression heteromeric α2β GlyR, gephyrin and NL2 in HEK293 cells. We performed unroofing to retain only the plasma membrane that is attached to the glass substrate and used total internal reflection fluorescence (TIRF) microscopy to limit excitation depth (Fig. 5F, see methods for details). Clearly, NL2, gephyrin and α2β GlyR forms micron-sized co-clusters at the plasma membrane. Since these clusters contain functional α2β GlyR, they are poised to generate electrical activities upon binding with neurotransmitter glycine. Local concentration of NRX upon binding with these clusters would signal the formation of cognate pre-synaptic specializations(*8*).

## Discussion

We show that only the ECDs of NL2 and NRX1β were resolved despite using full-length proteins, suggesting a flexible orientation between the TM and ECD. We identified structural mechanisms underlying the low affinity and non-cooperative binding between NL2 and NRX1β. Both full- and sub-stoichiometric NL2-NRX1β adhesion complexes are stable in solution. These complexes showed structural flexibilities that allow variations in the orientation between the two monomers of NL2 dimer. Despite low binding affinities, NL2 - NRX1β interaction alone is sufficient to tether lipid membranes in reconstituted proteoliposomes. Clustering of NL2 and NRX1β at inter-cellular junctions likely provide sufficient avidity to tether cellular membranes, and serves as signal to coordinate two contacting cells in generating functional synapse, through the recruitment of other essential synaptic factors.

The above findings raise a tempting mechanism of how NL2 – NRX1β interaction contributes to spatial and chemical precision in the formation of functional synapse (Fig 6). The plasma membranes of two cells come in proximity by chance and allow the formation of NL2—NRX1β adhesion complex (Fig. 6A). NL2 and NRX1β then cluster at the junction, providing more avidity. Flexible linkers between ECD and TM for both NL2 and NRX1β, combined with variable orientation between NL2 monomers make the adhesion complex amenable to membrane geometry (Fig. 6B, C). Clustering of NL2 and NRX1β also results in locally concentrating their intracellular domains, which are known to interact with synaptic scaffolding proteins that in turn recruit factors necessary for synaptogenesis(*8, 11, 26*). Since NL and NRX only cluster at the interaction site, this mechanism allows spatial coordination between two cells. The specificity between NLs and NRX, as well as between ICDs and respective scaffolding proteins, contribute to chemical specificity. NL2-NRX1β clustering likely initiates before synapse formation and serve as synaptogenesis signal, consistent with its ability in inducing synaptic formation in neurons(*41–43*)

**Figure 6.**
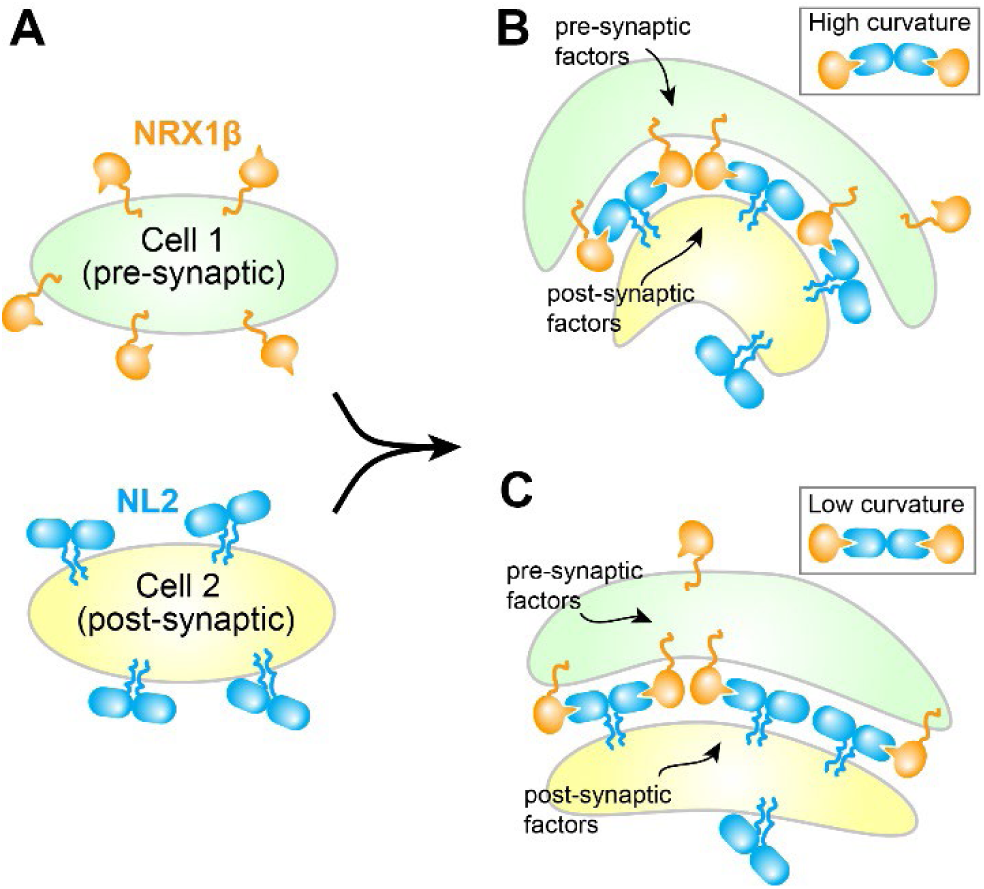
Illustration of NL2-NRX1 interaction at cellular junctions. (**A**) NL2 and NRX1 are randomly distributed on surface in the absence of inter-cellular contacts. ECD orientations with respect to membrane is flexible. Apo state NL2 adopts conformation that is incompatible with NRX1 binding. (**B**) and (**C**), NL2 and NRX1 cluster at cellular junctions, as full-or sub-stoichiometric complexes. Despite weak interactions, multiple NL2-NRX1 binding provide sufficient avidity for tethering two membranes. Conformational variations between monomers of NL2 dimer may provide flexibility of large clusters at (**B**) high and (**C**) low membrane curvatures.

The mechanism underlying NL2 NRX1β clustering is unclear. Since clusters are observed in HEK cells, they are unlikely to be dependent on any synapse-specific component. Clustering is less likely through additional ECD interactions previously proposed based on crystal contacts either(*31*), since such assembly implicates defined stoichiometry as well as observable interaction without crystallization, neither of which is supported by our observations. Instead, initial clustering/local concentrating may purely be a thermodynamic outcome -- NL2-NRX1β complex can only diffuse within the junction where membranes are sufficiently close, while NL2 or NRX1β alone do not have this limitation. This coincides with NRX1β being ∼ 3.6 folds more concentrated at junction and NL2 concentrating ∼ 10 ≈ 3.6^2^ folds, as NL2 has 2 binding sites (thus double the energy) but NRX1β has only 1. However, in cells, it might be much more complicated considering cross-interaction between other adhesion proteins and scaffolds, especially at cellular junctions where scaffolds in both cells may contribute to obstructed diffusion(*2, 26, 34*).

Tethering synaptic membranes through high avidity using multiple weaker interactions in clusters may offer advantages in regulation capability. For instance, NL2-NRX1β interaction can be easily displaced with competing factors, such as MDGA1(*32, 33*) that binds NLGN2 with nanomolar affinities. In addition, stable tethering most likely requires the involvement of other (SAM) proteins. Clearly, the properties and regulation of NL-NRX clusters at different stages of synaptogenesis requires further investigation to understand the elegant construction and regulation of synapses.

## Data availability

The density maps and structural model for the cryo-em data have been deposited with the Electern Microscopy Data Bank (EMDB) and the Protein Data Bank (PDB) with the following accession codes. NL2 apo (EMDB: EMD-29811, PDB: 8G7D); NL2-NRX1β conformation 0 (EMDB: EMD-29828, PDB: 8G7Y); NL2-NRX1β conformation 1 (EMDB: EMD-29829, PDB: 8G7Z); NL2-NRX1β conformation 2 (EMDB: EMD-29831, PDB: 8G81); NL2-NRX1β conformation 3 (EMDB: EMD-29830, PDB: 8G80).

## Acknowledgments

We thank Dr. Matthew Parker for the assistance in confocal microscopy. All members of the Wang laboratory for helpful discussions. Cryo-EM data was collected at the University of Texas Southwestern Medical Center Cryo-EM Facility, which is funded by the CPRIT Core Facility Support Award RP170644. This work is supported by NIH grant 1R35GM146860 and the McKnight Scholar Award to W.W. and by NIH grant R35GM119619 to K.J.

## Methods

### Cloning of NL2 NRX1β

The coding sequences of mouse NL2 and mouse NRX1β were amplified from pNICE-NL2(-) (Addgene item #15246) and pNICE-LAP-neurexin-1β (Addgene item #42575), respectively. They were subsequently cloned into Bacmam vectors for expression in Human embryonic kidney 293-T (HEK293T) cells(*44, 45*). The signal peptide of NL2 was replace with that of NL1 followed by an HA-tag (see Fig. S1). Both protein were constructed as a eGFP fusion at the C-terminus for affinity purification using anti-GFP antibodies(*46*).

### Protein Expression and Purification for NL2 Apo structure

Vector containing NL2 was transformed into DH10BacY competent cells (Geneva Biotech) to produce bacmids. Sf9 cells where transfected (Invitrogen) with the bacmids then amplified in Insect-XPRESS medium (Biowhitaker). Large scale protein expression was done using HEK293T cells grown in Freestyle 293 medium (GIBCO) containing 1% fetal bovine serum and 1% Penicillin-Streptomycin. Cells were infected with virus at a density of 2.0 x10^6^ cell/mL, induced using 10 mM sodium-butyrate 20 hours after infection, and grown for a total of 72 hours. After harvesting, cells were lysed using 10 mM HEPES-Na, 50 mM NaCl, 1 mM MgCl_2_ and 0.5 mM CaCl_2_. The protein was extracted using 0.5% LMNG in a 20 mM HEPES-Na, 200 mM NaCl, 1 mM MgCl_2_ and 0.5 mM CaCl_2_ buffer for 1 hour at room temperature. Protein was purified using GFP-nanobody resin and eluted by PPX digestion, then further purified using size exclusion chromatography using a Superose 6 (GE) column in a 20 mM HEPES-Na, 150 mM NaCl, 1 mM MgCl_2_, 0.5 mM CaCl_2_, 0.06% Digitonin, 0.05 mg/mL brain polar extract lipid buffer. Peak fractions were collected and concentrated.

### Protein Expression and Purification for NL2 NRX1β structures

Virus containing NRX1β was produced in like fashion to NL2. Large scale protein expression for each protein was done separately using the same method. HEK293T cells were infected with virus at a density of 2.0 x10^6^ cell/mL, induced using 10 mM Sodium-Butyrate 20 hours after infection, and grown for a total of 72 hours. After harvesting, cells were lysed using 10 mM HEPES-Na, 50 mM NaCl, 1 mM MgCl_2_ and 0.5 mM CaCl_2_. The protein was extracted using 0.5% LMNG in a 20 mM HEPES-Na, 200 mM NaCl, 1 mM MgCl_2_ and 0.5 mM CaCl_2_ buffer for 1 hour at room temperature. Protein was purified using GFP-nanobody resin and eluted by PPX digestion, then further purified using size exclusion chromatography using a Superose 6 (GE) column in a 20 mM HEPES-Na, 150 mM NaCl, 1 mM MgCl_2_, 0.5 mM CaCl_2_, 0.05% LMNG buffer. Peak fractions where collected and concentrated.

### CryoEM Sample Preparation

NL2 and NRX1β were purified separately and concentrated down. They were then mixed at a one to one molar ratio for a final OD of 3.1, and incubated on ice for 10 minutes. Samples were applied to a glow - discharged Quantifoil R1.2/1.3 400-mesh gold holey carbon grid (Quantifoil, Micro Tools GmbH, Germany), blotted under 100% humidity at 4°C and vitrified in liquid ethane using a Mark IV Vitrobot (FEI).

### CryoEM data acquisition and processing

Micrographs were collected using a Titan Krios microscope (Thermo Fisher) with a K3 Summit direct electron detector (Gatan) operating at 300 kV. SerialEM was used for automated data acquisition with GIF-Quantum energy filter set to a slit width of 20 eV. Images were recorded in the super-resolution counting mode with the pixel size of 0.415 Å. Micrographs were dose-fractioned into 50 frames with a dose rate of 1.8 e^-^/Å/frame for NL2 apo and 1.4 e^-^/Å/frame for NL2-NRX1β complex. 4988 and 6615 movies were collected for NL2 apo and NL2-NRX1β complex, respectively.

Motioncorr2(*47*) was used for Fourier truncation (2-fold binning, 0.83 Å pixel size afterwards), motion correction and dose weighting of the movie frames, with subsequent CTF correction performed using CTFFIND 4 program(*48*). Image processing steps were carried out in RELION 3(*49*) and cryoSparc(*50, 51*), as illustrated in Fig. S2. Initially particle picking using the Laplacian-of-Gaussian blobs allowed 2D classification to obtain good class-averages, which were used for templates for reference-based autopicking. Resulting particles were cleaned up with further rounds of 2D classifications. Good 2D classes were used to generate initial 3D model and subsequent 3D classification and refinement as detailed below.

For NL2 apo, 3D heterogeneous refinement with 3 classes in cryoSparc(*50*) resulted in one good class, containing 77, 577 particles. This class was further subjected to Non-Uniform refinement(*51*) and reached an overall resolution of ∼3.3 Å.

For NL2-NRX1β complex, 3D classification into 8 classes resulted in 2 good classes, class 2 and class 5, with class 2 having inverted projection. After flipping Z for class 2 using Chimera software(*52*) and further 3D classification for both class 2 and class 5 (Fig. S2I), 1 sub-stoichiometric (1 NRX1β: 1 NL2 dimer, Conf. 0), and 3 full-stoichiometric complex (2 NRX1β: 1 NL2 dimer, Conf. 1-3) complex structures were identified. Particles of each structure was CTF-refined and Bayesian polished in RELION after Non-Uniform refinement in cryoSparc. A final round of Non-Uniform refined with polished particles results in the final density maps with overall resolutions of 3.9 Å for Conf. 0 and 3.2 Å for Conf. 1-3 (Fig. S2J-L). Good local resolutions extending beyond 2.5 Å were observed in NL2, while those of NRX1β are more moderate, especially in parts further away from NL2 (Fig. S2M). Resolutions were estimated by applying a soft mask around the protein densities with the Fourier Shell Correlation (FCS) 0.143 criterion. Local resolutions were calculated using Resmap(*53*).

### Model building and refinement

The model of apo NL2 was made by aligning the NL2 structure (PDB ID: 3BL8) to the density map using Chimera(*52*). The model was then adjusted and the amino acids were mutated manually using Coot(*54*). The model was refined using real space refinement program in PHENIX(*55*). The final NL2 model begins at R40 and shows all amino acid except the following due to low map density: chain A-E151-S174, D553-N563, and ends at L608, chain B-E151-s174, I557-P562, and ends at N610. For the NL2-NRX1β complexes, each monomer of atomic model (PDB ID: 3BIW) was fitted into experimental maps using Chimera, manually adjusted using Coot, and refined in PHENIX (see supplementary Table 1 for statistics) similarly to the NL2 Apo model. Statistics of cryo-EM data processing can be found in TableS1. For the configuration 0 model the NL2 subunit begins at E39 for chain A and R40 for Chain B and shows all amino acids except for as follows due to low map density: chain A D152-G175, F557-K561, and ends at N610, chain B D152-G175, F557-K561, and ends at H 609. The NRX1β subunit begins at A83 and shows all amino acids except for as follows due to low map density: E162-I168, R202-L231, and ends at G289. For configuration 1 both NL2 chains start at R40 and end at L608, with no modeling for E151-D173, T555-N563 for chain A, and E151-D173, D554-N563 for chain B. For configuration 2 both NL2 chains start at R40 and end at L608, with no building for E151-D173 and P552-P562 for chain A, and E151-D173 and K556-P562 for chain B. For Configuration 3 model the NL2 subunit begins at R40 for both chains A and B. Omissions due to low density for the NL2 subunit are as follows: T150-S174 for chain A, T150-D173 for chain B, D554-N563 for both chains, and both chains end at L608. The NRX1β subunit chains for configuration 1, 2, and 3 all have the same missing amino acids, with the chains beginning at A83 and missing R202-L231, and ending at L289.

### mPEG glass preparation

Glass coverslips were cleaned by soaking in 100 mL of Piranha solution for 90 minutes, sonicating for 10 seconds and the start and every 45 minutes after. The slides were rinsed with MiliQ H_2_O 5 times. The cleaned slides were etched using 1 M KOH and soaked for 30 minutes sonicating for 1 minuet at the start and every 15 minutes after. The coverslips were then rinsed and dried using filtered air. The coverslips were then preheated to 90°C and 10 µL of 25% mPEG-sliane 5k with 1% Biotin mPEG-silane 5k in DMSO was added to the coverslip. A second coverslip was added to coat the whole area. The coverslips were then incubated at 90°C for 30 minutes, rinsed 5 times with MiliQ H2O, and dried using filtered air. And stored at −20°C for later use.

### Proteoliposome reconstitution and TIRF Imaging

A 90:10:0.1:0.1 1-palmitoyl-2-oleoyl-glycero-3-phosphocholine (POPC): 1-palmitoyl-2-oleoyl-sn-glycero-3-phospho-L-serine (POPS): 1,2-dioleoyl-sn-glycero-3-phosphoethanolamine-N-(cap biotinyl) (Biotin-PE):1,2-dioleoyl-sn-glycero-3-phosphoethanolamine-N-(7-nitro-2-1,3-benzoxadiazol-4-yl)(NBD-PE) (Avanti) lipid mixture was dried using Argon gas, and placed in a vacuum desiccator overnight. After drying lipids were resuspended in miliQ H_2_O at 20 mg/ml and sonicated until clear, then solubilized with 0.5% DDM. Lipids were added to a purified NL2 protein at a ratio of 1:10 protein to lipid and diluted down with a 40 mM HEPES-Na, 300 mM NaCl buffer to a final concentration of 20 mg/ml. Bio-Beads SM-2 (BIO-RAD) where added to the mixture to remove detergent, and the mixture was rotated at 4°C. The Bio-Beads were changed 2X every 3 hours and change 1X overnight. For NRX1β a second lipid mixture was made in similar manner with the composition being 90:10:0.1:0.1 POPC:POPS:Biotin-PE: 1,2-dioleoyl-sn-glycero-3-phosphoethanolamine-N-(lissamine rhodamine B sulfonyl) (Rhod-PE), and the protein was reconstituted for a ratio of 1:20 protein to lipid. The reconstituted proteins were mixed together at a concentration of 0.1 mg/mL and incubated for 30 minutes, then further diluted for a final concentration of 0.33 ng/mL using 2 mg/mL BSA 20 mM HEPES-Na, 150 mM NaCl (dilution buffer). To the clean slides 15 µL of 280 nM NeutrAvidin in 5 mg/mL BSA, 20 mM HEPES-Na, and 150 mM NaCl was added to each coverslip and incubated at room temp for 15 minutes. They were then washed with 1 mL five time with 20 mM HEPES-Na 150 mM NaCl.

Then 15 µL of the reconstituted vesicles were added to the coverslip and incubated at room temp for 15 minutes, then rinsed with the dilution buffer 9 times. The slides were then imaged using total internal reflection illumination. This imaging was achieved using an in-house built prism-type system using the Gem 488 laser (Laser Quantum) and the Gem 560 nm laser (Laser Quantum) and components from Thorlabs at 10% power with an exposure time of 200 ms. A Leica DM6 FS microscope equipped with a 60x 1.2NA water immersion objective and a Hamamatsu flash 4.0 V3 camera was used for imaging, images were collected using a PC running Metamorph (Molecular Devices).

### Vesicle detection and colocalization analysis

Imaged vesicles were detected and localized using the “gaussian mixture-model fitting” algorithm in u-track(*56*) (Version 2.2.1; https://github.com/DanuserLab/u-track), using the iterative mixture-model fitting option. For both channels, the Gaussian standard deviation was taken as 1.1 pixel (113 nm), and the α-values for the residuals test, amplitude test and distance test were respectively set to 0.05, 0.1 and 0.05. The α-value for local maxima detection was set to 0.05 for the 560 channel (NRX1β) and 0.1 for the 488 channel (NL2). Visual inspection confirmed that these detection parameters detected all objects, both dim and bright, with minimal false positives or false negatives.

Object-based colocalization analysis was then performed on the detected vesicles, based on the nearest neighbor distances between detected objects in the two channels, as described(*36*), using a colocalization radius of 3 pixels (code available at https://github.com/kjaqaman/conditionalColoc). Colocalization analysis was performed on either all vesicles (all circles in Fig. S4A), or on those with intensities in the top 75% (green, cyan and magenta circles in Fig. S4A), in the top 50% (green and cyan circles in Fig. S4A) or in the top 25% (green circles in Fig. S4A). To assess the significance of any measured colocalization, i.e. to assess how unlikely it was to have arisen by chance, a “nullTR” colocalization measure (*computational null control*) was calculated for each sample, where the objects in one of the two channels were replaced by points on a grid. The real data colocalization measure was then compared to the nullTR measure using a Wilkoxon rank-sum test to compare medians, yielding the reported p-values.

### Cell imaging using confocal microscopy

Two plates HEK293T cells were transfected separately with NL2-eGFP and NRX1β - mCherry or NL2-mCherry and NRX1β-eGFP using a Lipofectamine 3000 Transfection Kit (Invitrogen) and grown overnight. The cells where then mixed together on a collagen treated glass-bottom dish (Mattek) and allowed to grow for 3 hours at 37°C. The cells were then fixed using a 4% Paraformaldehyde (Electron Microscopy Sciences) in DPBS (gibco) for 15 minutes at room temperature. After fixing the cells where imaged using a Nikon Ti2E microscope equipped with a Yokogawa CSU X1 spinning disk using a 60× oil immersion objective with the appropriate filter set and images collected with 488 nm and 561nm laser at 20% power. Images were processed using imageJ (see below).

### Quantification of fluorescence intensity

Intensity analysis was preformed using imageJ. The background of the image was determined at areas with no cell, and subtracted from measurements. Using the Plot Profile function on imageJ we plotted the junction fluorescence intensity of the two cells in the eGFP channel and used 1/3^rd^ of the measured peak value as the threshold for region of interest (ROI), within which mean fluorescence intensities were calculated. For the same cell, we used similar method to quantify fluorescence intensity of the non-junction membrane. The junction intensity was the divided by the non-junction intensity to give us the fold difference. To estimate protein concentration, the fluorescence intensities were measured at different concentrations (0, 0.1 µm, 1 µM, 3 µM, 9 µM) of soluble eGFP proteins. Number of molecules contributing to fluorescence were calculated using *N = [GFP] x V_c_ x N_A_*, where *V_c_* is the effective confocal volume and *N_A_* is the Avogadro’s number (6.02 x 10^23^). *V_c_* is estimated using 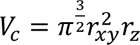, where 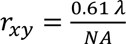 and 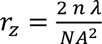 are the diffraction-limited e^-2^ radius in x-y and z directions, respectively.*λ* (excitation wavelength) is 488 nm, *n* (sample refractive index) is 1.33 and *NA* (objective numerical aperture) is 1.4.

### Purification of NRX ECD

NRX1β ECD his-tag purification was performed by transforming the NRX1β ECD (aa 81 - 311) DNA into BL21 competent *E.coli* cells and grown into a large scale 1 L prep. The cells where grown to a density of 0.8∼1.0 OD and induced with 0.2 mM IPTG then grown at 16°C for 24 hours. Once the cells where harvested they were resuspended into binding buffer (20 mM Tris, pH8.0, 500 mM NaCl) then sonicated for 1 minute 3 times with stirring at 4°C for 5 minutes in between each sonication. The cells were then centrifuged and the supernatant was loaded on to Ni-NTA resin (Biosciences). The protein bound resin was washed with 50 mL of binding buffer, then washed with 50 mL of 20 mM Tris, pH8.0, 500 mM NaCl, 40 mM Imidazole. The protein was then eluted off the resin using 50 mL of 20 mM Tris, pH8.0, 500 mM NaCl, 500 mM Imidazole. The protein was concentrated and further purified using size exclusion chromatography using an ENrich Sec 70 10X300 Column (BioRad).

### Fluorescent labeling of NRX ECD

0.5mg of NRX1β ECD was diluted in labeling buffer (20 mM HEPES 150 mM NaCl, 0.05% LMNG) and fluorescently labeled with 200 mM of Alexa Fluorophore 488-NHS by overnight incubation with. The protein dye mixture was then desalted in a pre-packed Sephadex G-25M column (PD-10, GE Healthcare) using labeling buffer. Labeled protein was concentrated and further purified using size exclusion chromatography using the SEC 70 column in the labeling buffer.

### Fluorescent Polarization

Fluorescent polarization was measured for NRX1β ECD labeled with Alexa Fluorophore 488, during titration of either NL2 WT or NL2mut (H279Y G282K). Using a 96 well Optical Bottom PolymerBase plate (ThermoFisher) a triplicate was performed for each NL2 WT and NL2mut. The titration was performed in 20 mM HEPES, 150 mM NaCl, 1 mM MgCl_2_, 0.5 mM CaCl_2_ with 8 nM NRX1β ECD – AF488. The NL2/NL2mut concentrations were: 15 nM, 30 nM, 62.5 nM, 125 nM, 250 nM, 500 nM, 1 µM, 2 µM, 4 µM, and 8 µM. The plate was imaged using Spark Multimode Microplate Reader (TECAN) Fluorescent polarization methods with an excitation wavelength of 490 and an emission wavelength of 540.

### Co-Transfection and TIRF Imaging of Clusters

22 mm x22 mm #1 Slip-rite cover glass (Thermo) was cleaned using Sulfuric Acid for 30 minutes, then rinsed with Mili-Q water 3 times and dried in a sterile hood for one hour. The cover slips where then placed into a 35 mm dish and coated with 100 µg/mL of collagen and incubated for one hour and 37°C. They were then rinsed with Phosphate-Buffered Saline (PBS) three times and dried in sterile hood for one hour. The plates containing the coverslips were then seeded with 0.4 million of HEK293T cells and transfected with NL2 containing a Halo tag on the C-term, GPHN (residues 2-736 with N-terminal mCherry tag), GlyRα2em, and GlyRβem at a ratio of 2:2:1:1 using a Lipofectamine 3000 Transfection Kit (Invitrogen) and grown for 20 hours(2). GlyRα2em, and GlyRβem has been described before (*45*). Briefly, the GlyRα2em subunit has a GSSG substitution at residues Q317-K381; the GlyRβem has two modifications; residue N334-N377 was replaced by GGSSAAA-monomeric enhanced <u>green fluorescent</u> <u>protein</u> (mEGFP)-SGSGSG, and at the N terminus a PA-tag (GVAMPGAEDDVV) (*57*) and PreScission protease site (LEVLFQ/GP)(*58*) were inserted. The cells were then washed at room temperature 3 times with PBS, then sonicated at 20% power for 1 second using PBS supplemented with 2% paraformaldehyde (PFA). After sonication the cells were washed with PBS and incubated at room temperature for 15 minutes in PBS containing 4% PFA. The cells were then washed with PBS 3 times for 5 minutes each. To label the NL2 the cells where blocked for 15 minutes using PBS with 1% bovine serum albumin (BSA), then the primary antibody, HA-Tag C29F3 rabbit mAb (Cell Signaling) was added at a ratio of 1:2000 and the cells were incubated for one hour at room temperature. The cells were then rinsed with PBS and the secondary antibody, Goat Anti-Rabbit IgG, Highly Cross-Adsorbed, Labeled with CF660C dye (BIOTIUM), was added to the cells at a concentration of 1 µg/mL and incubated at room temperature for one hour. The cells were then rinsed once more with PBS, and the cover slip was then imaged using the same TIRF imaging methods as described above.

