## Supplemental data for "Weaker neuroligin 2 – neurexin 1β interaction tethers membranes and signal synaptogenesis through clustering"

### Supplemental information

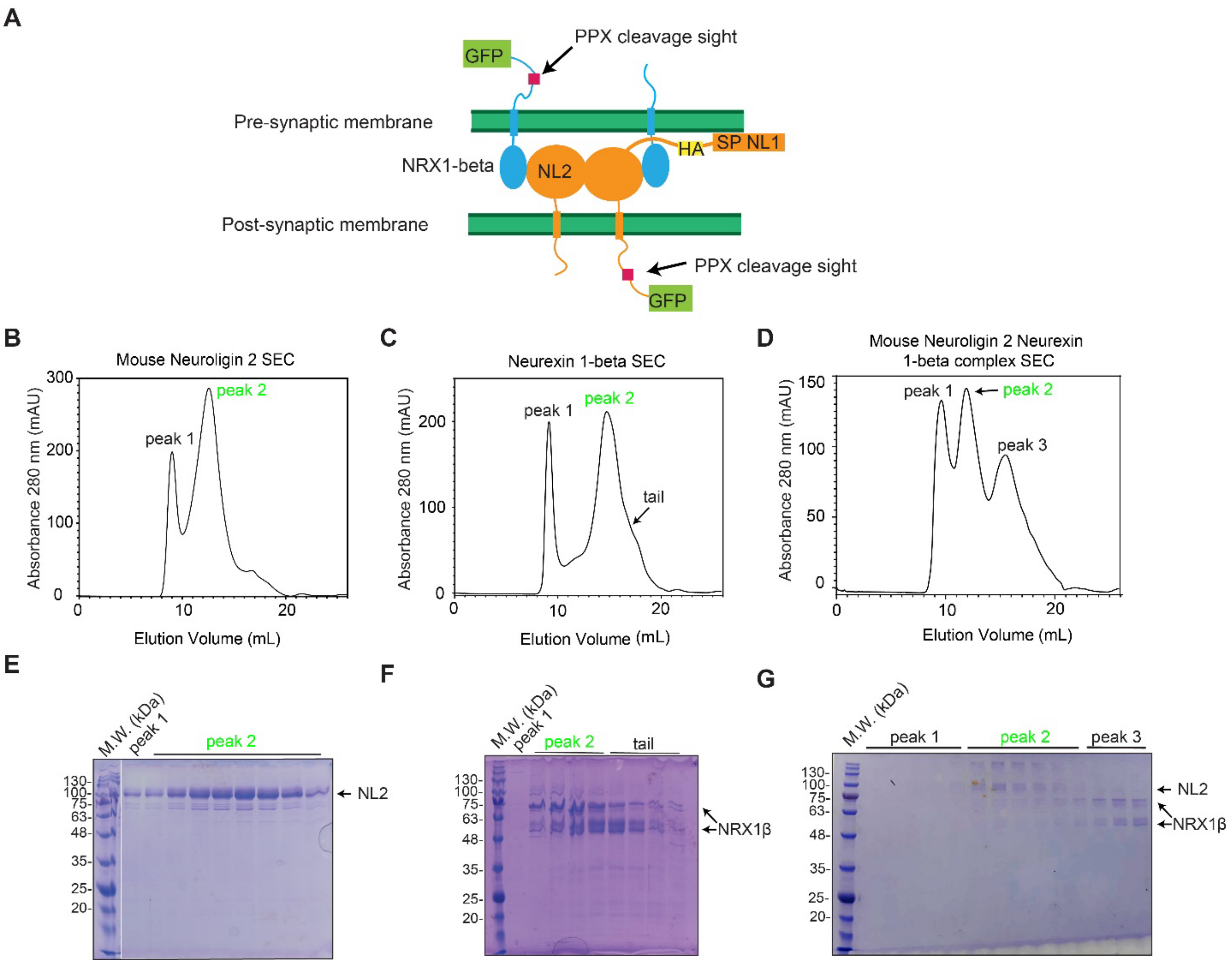


**Figure S1. Expression and purification of NL2 and NRX1β**. (A) Illustration of full-length NL2 and NRX1β, with signal sequence, tags, and precision protease (PPX) cleavage sites shown. GFP tag allowed purification using GFP-nanobody resin. (B-D) SEC elution profiles of respective proteins, with peak 1 representing the void volume, and peak 2 being the desired protein/complex. (E-G) SDS PAGE of respective SEC fractions.

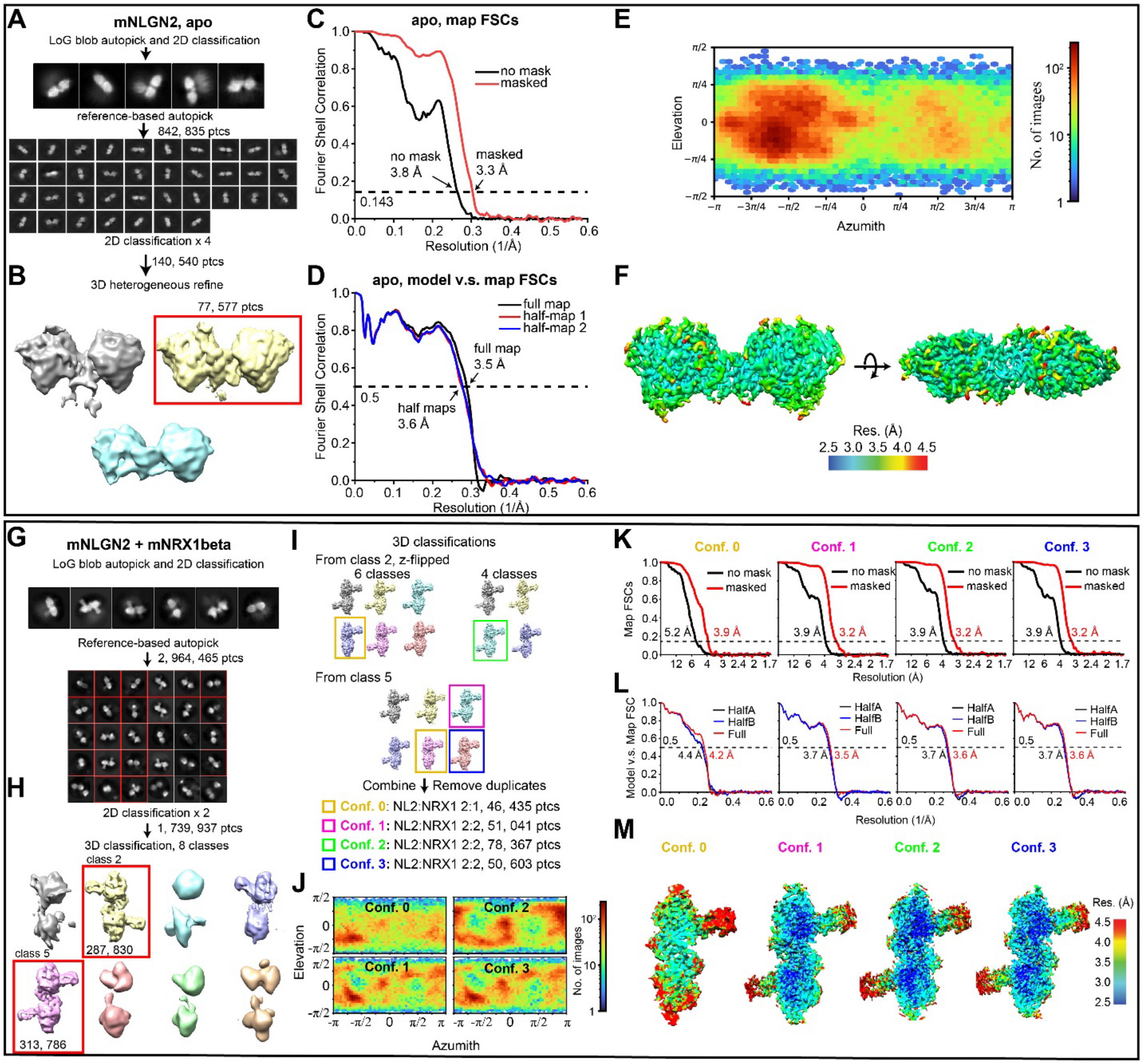


**Figure S2**. **Overview of cryo-EM data processing. (A-F**) Processing of NL2 apo dataset. (**A**) Particle picking and 2D classification. (**B**) Classification into three 3D classes results in one good class containing 77, 577 particles. Fourier-Shell Correlations (FSC) between (**C**) two half maps and (**D**) atomic model and maps, with resolution at FSC of 0.143 (between maps) and 0.5 (model v.s. maps) indicated. (**E**) Angular distribution of particles in final refinement. (**F**) Density map colored according to local resolutions view from side and top. **(G-M**) Processing of mNL2-

mNRX1β dataset. (**G**) Particle picking and 2D classification. Red rectangles indicate selected classes. (**H**) Classification into eight 3D classes results in two good class (red rectangles) with the numbers of particles in each class indicated. (**I**) Further 3D classification yielded good resolution maps with distinct conformations (yellow, purple, green and blue respectively corresponds to Conf. 0, 1, 2, 3). (**J**) Angular distributions of particles in final refinement for each conformation. FSCs between (**K**) two half maps and (**L**) atomic model and maps, with resolutions at 0.143 FSC (between maps) and 0.5 FSC (model v.s. maps) indicated. (**M**) Density maps sliced through center, viewed from top and colored according to local resolutions. Local resolutions were estimated using ResMap 1.1.4.

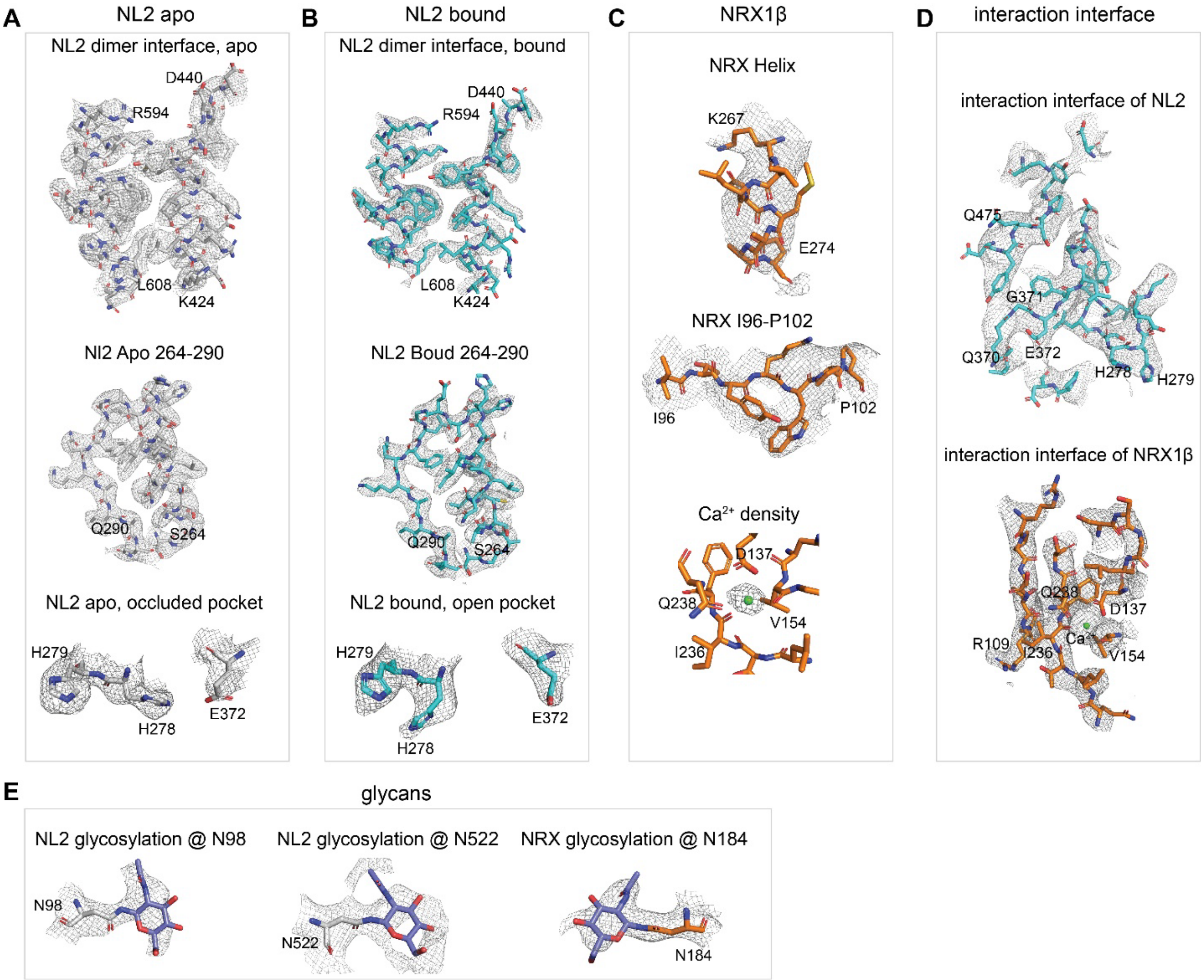


**Figure S3. Density maps for representative regions**. Regions of (**A**) NL2 apo (gray), (**B**) NL2 bound (cyan), (**C**) NRX1β (orange), (**D**) NL2-NRX1β interaction interface and (**E**) N-linked glycans (light blue) are shown.

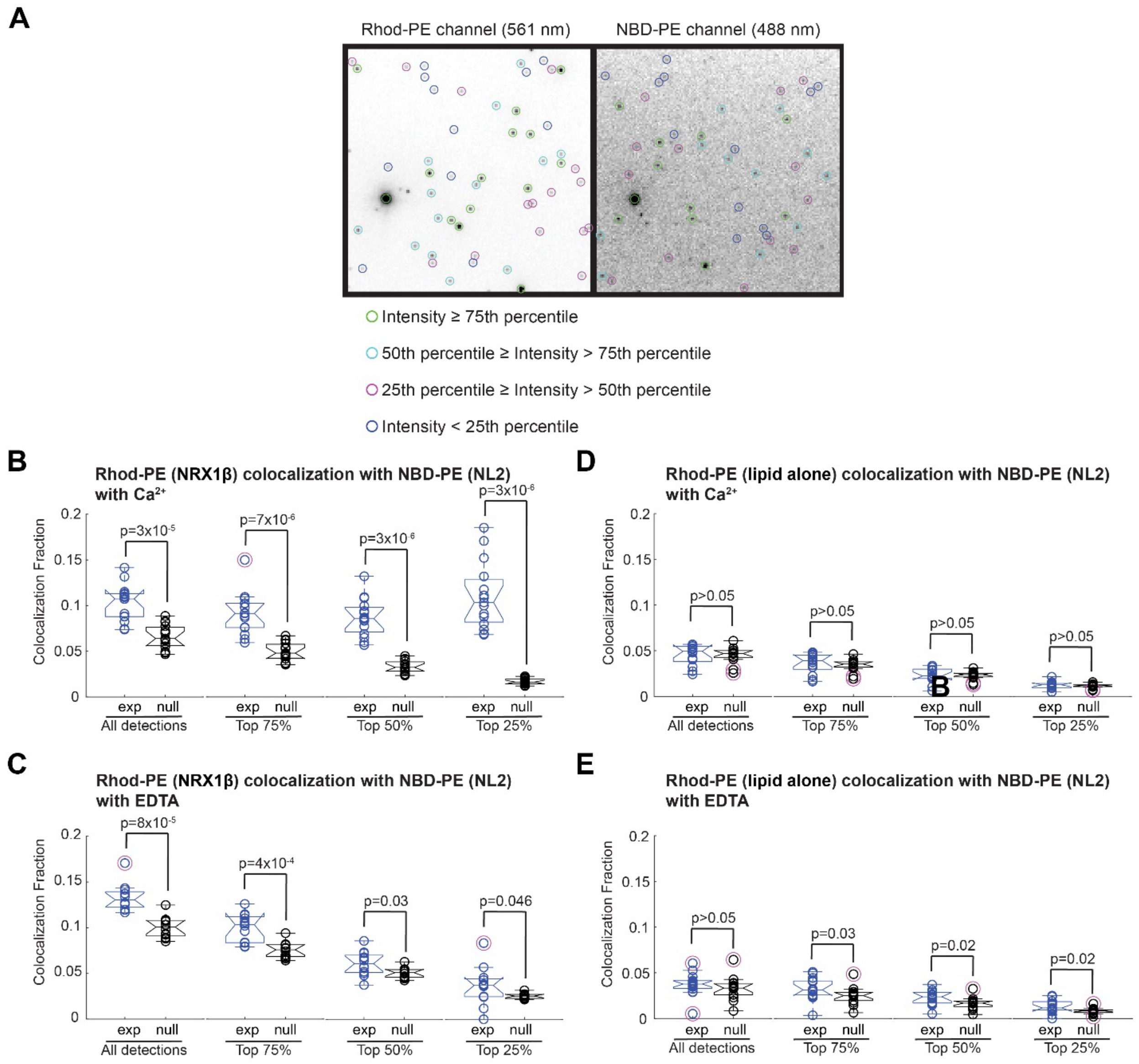


**Figure S4. Object-based colocalization analysis**. (**A**) Illustration of detections with object intensities falling into indicated ranges. (**B** and **C**) Object-based co-localization analysis of NRX1β (Rhod-PE) with NL2 (NBD-PE) vesicles in the presence of (**B**) $Ca^{2+}$ and (**C**) EDTA. (**D** and **E**) Co-localization analysis of lipid alone (Rhod-PE) with NL2 (NBD-PE) vesicles in the presence of (**D**) $Ca^{2+}$ and (**E**) EDTA. In B-E, for data presentation description and number of data points used in analysis, see Fig. 5.

### Supplementary Table 1 Cryo-EM data collection, refinement and validation statistic

| | Mouse Neuroligin 2 Dimer | Mouse Neuroligin 2 Dimer, one Nerexin1-beta | Mouse Neuroligin 2 Dimer, two Nerexin1-beta (Configuration One) | Mouse Neuroligin 2 Dimer, two Nerexin1-beta (Configuration Two) | Mouse Neuroligin 2 Dimer, two Nerexin1-beta (Configuration Three) |
|---|---|---|---|---|---|
| **Data collection and processing** | | | | | |
| Magnification | 105,000 | 105,000 | 105,000 | 105,000 | 105,000 |
| Voltage (kV) | 300 | 300 | 300 | 300 | 300 |
| Electron exposure (e–/Å$^2$) | 90 | 70 | 70 | 70 | 70 |
| Defocus range (μm) | -1.0 to -2.5 | -1.0 to -2.5 | -1.0 to -2.5 | -1.0 to -2.5 | -1.0 to -2.5 |
| Pixel size (Å) | 0.83 | 0.83 | 0.83 | 0.83 | 0.83 |
| Symmetry imposed | *C1* | *C1* | *C1* | *C1* | *C1* |
| Initial particles(no.) | 674,761 | 2,964,465 | 2,964,465 | 2,964,465 | 2,964,465 |
| Final particles (no.) | 77,577 | 46,435 | 51,041 | 78,367 | 50,603 |
| Map resolution (Å) | 3.28 | 3.92 | 3.22 | 3.22 | 3.25 |
| FSC threshold | 0.143 | 0.143 | 0.143 | 0.143 | 0.143 |
| **Refinement** | | | | | |
| Initial model used (PDB code) | 3BL8 | 3BIW | 3BIW | 3BIW | 3BIW |
| Model resolution (Å) | 3.28 | 3.92 | 3.22 | 3.22 | 3.25 |
| FSC threshold | 0.5 | 0.5 | 0.5 | 0.5 | 0.5 |
| Model composition | | | | | |
| Non-hydrogen atoms | 8346 | 19146 | 11094 | 11148 | 11067 |
| Protein residues | 1076 | 1258 | 1427 | 1432 | 1425 |
| Ligands | NAG:4 | NAG:5 CA:1 | NAG:6 CA:2 | NAG:6 CA:2 | NAG:6 CA:2 |
| *B* factors (Å$^2$) | | | | | |
| Protein | 77.52 | 157.02 | 88.21 | 120.42 | 74.30 |
| Ligand | 136.32 | 186.97 | 128.38 | 166.43 | 142.03 |
| R.m.s. deviations | | | | | |
| Bond lengths (Å) | 0.003 | 0.003 | 0.004 | 0.003 | 0.002 |
| Bond angles (°) | 0.516 | 0.542 | 0.582 | 0.563 | 0.502 |
| Validation | | | | | |
| MolProbity score | 1.77 | 1.98 | 1.90 | 1.76 | 1.78 |
| Clashscore | 6.07 | 10.60 | 8.99 | 8.31 | 8.55 |
| Rotamer outliers (%) | 0 | 0 | 0 | 0 | 0 |
| Ramachandran plot | | | | | |
| Favored (%) | 93.33 | 93.32 | 93.62 | 95.55 | 95.39 |
| Allowed (%) | 6.67 | 6.60 | 6.38 | 4.45 | 4.61 |
| Outliers (%) | 0.0 | 0.0 | 0.0 | 0.0 | 0.0 |
